# Human Cytomegalovirus Intrahost Evolution–A New Avenue for Understanding and Controlling Herpesvirus Infections

**DOI:** 10.1101/009571

**Authors:** Nicholas Renzette, Laura Gibson, Jeffrey D. Jensen, Timothy F. Kowalik

**Affiliations:** Department of Microbiology and Physiological Systems, University of Massachusetts Medical School, Worcester MA 01655, USA; Departments of Pediatrics and Medicine, Divisions of Infectious Diseases and Immunology, University of Massachusetts Medical School, Worcester MA 01655, USA; Program in Bioinformatics and Integrative Biology, University of Massachusetts Medical School, Worcester MA 01655, USA; School of Life Sciences, Ecole Polytechnique Fédérale de Lausanne (EPFL), Lausanne 1015, Switzerland; Swiss Institute of Bioinformatics (SIB), Lausanne 1015, Switzerland; Immunology and Microbiology Program, University of Massachusetts Medical School, Worcester MA 01655, USA

## Abstract

Human cytomegalovirus (HCMV) is exquisitely adapted to the human host, and much research has focused on its evolution over long timescales spanning millennia. Here, we review recent data exploring the evolution of the virus on much shorter timescales, on the order of days or months. We describe the intrahost genetic diversity of the virus isolated from humans, and how this diversity contributes to HCMV spatiotemporal evolution. We propose mechanisms to explain the high levels of intrahost diversity and discuss how this new information may shed light on HCMV infection and pathogenesis.

## Introduction

Collectively, the cytomegaloviruses are ancient viruses that appear to infect the majority of vertebrates, but each particular cytomegalovirus infects a single host species with little evidence that the viruses jump between hosts. These traits have contributed to cytomegaloviruses adapting over long time scales (e.g. millennia) to the particular ecological niches in which they occupy. In particular, human cytomegalovirus is a β-herpesvirus that produces lifelong infections in most humans. In most individuals, HCMV infections result in a mild febrile illness, but can lead to severe symptoms in the immune-compromised or neonates. Nearly all HCMV infections result in widespread dissemination throughout the body, with diverse cell types such as epithelial, endothelial, fibroblast, and smooth muscle cells supporting productive viral infection. Further, the virus induces a myriad of immunomodulatory pathways to subvert the host innate and adaptive response [1]. Many excellent studies and reviews (for example, see [2] or [3]) have focused on the long term evolution of the virus and have highlighted the molecular characteristics that have helped create a highly successful pathogen. However, work mostly from the past decade has shown that the virus evolves on much shorter timescales, on the order of days or months, and within individual infected hosts. The purpose of this review is to discuss recent findings illustrating the complexity of short-term HCMV evolution and how it may relate to clinical manifestations of infection.

## Diversity of Human Cytomegalovirus in Human Hosts

Work published more than 30 years ago based on restriction length polymorphisms (RFLP) has shown that HCMV exhibits unexpected levels of genetic diversity between individuals (i.e., *inter*host variability) [4]. These results were later confirmed with targeted re-sequencing of genetic loci, primarily focusing on glycoproteins such as gB, gN and g0, as well as whole genome data from both low passage isolates and clinical samples [5,6]. However, work reported more recently has shown that HCMV also exhibits significant levels of genetic diversity within a single individual (i.e., *intra*host diversity). Similar to interhost variability, most intrahost diversity data has focused on the HCMV glycoproteins. Various assays, such as PCR-RFLP [7], DNA sequencing [8], single strand conformation polymorphism [9] and heteroduplex mobility analysis [9], show that these regions can be variable within individuals. However, these studies were limited by the technologies available, and suffered from issues associated with low sampling depth and narrow breadth of the HCMV genome. Nevertheless, they regularly showed that mixed infections accounted for roughly one quarter to one half of HCMV infections over a wide range of human populations, including infants with congenital infection [10], people with HIV/AIDS [11,12], transplant recipients, and immunocompetent children [13] and adults [14]. An important paper from 2011 began to show the full extent of HCMV intrahost diversity by using ultra-deep pyrosequencing to study the gO, gN and gH loci from HCMV-infected transplant recipients [15]. They found that all patients studied had mixed infections, with as many as 6 genotypes observed in a single patient. Similarly, deep sequencing of 81,224 base pairs of the HCMV genome directly from a glioblastoma multiforme (GBM) tumor revealed high genetic diversity of the virus including an overabundance of hypervariable loci, in which all four nucleotides were observed at a single position in the viral genome. Work that combined genomics with high throughput sequencing showed that HCMV diversity was not limited to a subset of loci, but rather spanned the entire genome [16]. In fact, nearly every open reading frame (ORF) of the HCMV genome exhibited a measurable level of intrahost genetic diversity. Importantly, HCMV diversity was described quantitatively using various metrics and shown to rival that of some RNA viruses, the benchmark of highly diverse viral populations (Figure 1). From these studies, it has been become clear that HCMV is highly diverse within humans.

**Figure 1:**
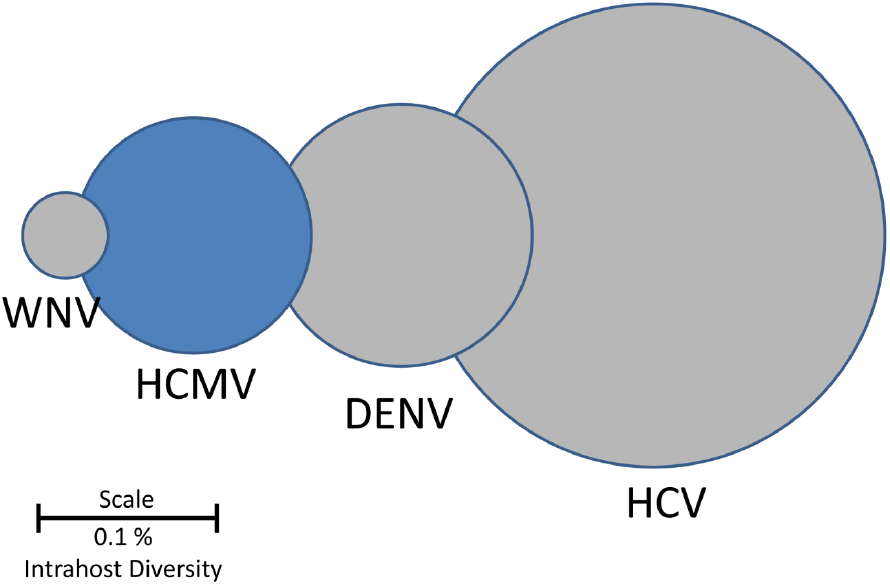
HCMV Intrahost genetic diversity as compared to RNA viruses. Viral intrahost diversities are represented as circles with the diameters, drawn on a log scale, representing reported values of diversity. The genetic diversity of HCMV populations is comparable to those of RNA viruses, an unexpected result given that HCMV is a large dsDNA virus. Values were obtained from [16] and references therein. Abbreviations are as follows: WNV: West Nile Virus, HCMV: human cytomegalovirus, DENV: dengue virus, HCV: hepatitis C virus. Scale bar represents the diameter of a genetic diversity value of 0.1%.

## Mechanisms of HCMV Diversity

Given the high levels of HCMV intrahost diversity, examination of its source is warranted. Currently, there is no clear mechanism to explain the observed diversity, though the literature does offer several non-mutually exclusive mechanisms that could be playing a role. High levels of replication during primary infection could contribute to the generation of *de novo* mutations in each host, though the total number of mutations generated may be limited by the proofreading activity of the viral DNA polymerase [17]. The excess of low frequency mutations in HCMV intrahost populations [15] as well as the star phylogeny of clones sampled from the populations [16] are consistent with this pattern (Figure 2A). However, clones from HCMV clinical samples also reveal that some members of the population are highly divergent from the most common sequence, a result consistent with reinfection (Figure 2B). HCMV reinfections appear to be common in both immunocompromised and healthy individuals [4,18-20]. Indeed, reinfections in some populations may occur at an annualized rate of 10%, comparable to the annualized rate of new infections [21]. Thus, reinfections offer a route for repeatedly introducing diversity into HCMV intrahost populations. However, this mechanism does not immediately appear to address the diversity observed in very young patients, specifically congenitally infected neonates. Here, the explanation may lie in the transplacental route of transmission associated with congenital infection. One recent study showed that maternal-to-fetal transmission may represent 10s to 100s of unique HCMV virions [22]. In contrast, transmission associated with RNA viruses is associated with a much lower number of virions, with bottlenecks so restrictive that only a single sequence transmits as in the case of HIV [23,24]. In this extreme case of a single virion transmission, all pre-existing viral population diversity is lost and must be generated *de novo* during replication in the newly infected host. More relaxed bottlenecks would result in 100s of HCMV virions being transmitted during congenital infections. In this circumstance, a portion of the pre-existing diversity present in the mother may be transmitted to the fetus. Whether this mechanism contributes to HCMV diversity in other transmission contexts, such as in healthy children or adults, is unknown.

**Figure 2:**
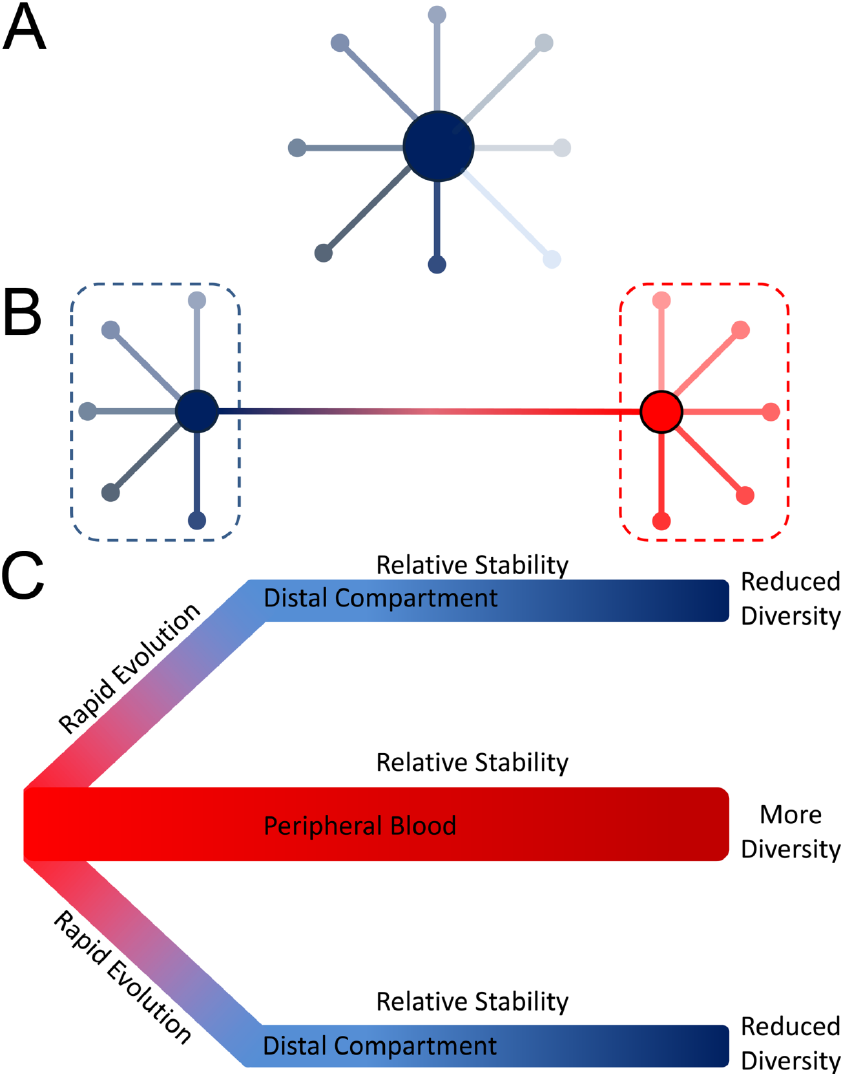
Models of HCMV Diversity and Evolution. **Panel A** shows an example of the structure of an HCMV intrahost population. Minor variants in the population are distributed around a central sequence. Most variants are only a few mutational steps away from this central sequence. This distribution of sequences is known as a star phylogeny and can be formed by the high levels of replication associated with primary infection, where most de novo mutations will be rare and shared by few members of the population. **Panel B** shows a more complicated population structure which could result from reinfection. In this example, the blue and red dashed lines represent strains, with minor variants radiating from the central sequence of each strain. The genetic distance between strains is significantly greater than the genetic distance of minor variants from their respective central sequences. **Panel C** depicts a model of HCMV evolution derived from congenital infection data. In this model, the peripheral blood harbors the most diverse sub-population of HCMV within the body. During dissemination to distal compartments, the viral population can rapidly evolve, either due to natural selection or stochastic mechanisms such as population bottlenecks, leading to populations in the distal tissues or organs being genetically differentiated from the peripheral blood compartment. This phenomenon is known as compartmentalization. The populations in the distal compartments are also less genetically diverse than those of the peripheral blood. Once HCMV has disseminated to compartments or if the virus remains in the peripheral blood, the populations become relatively stable and many mutations remain at similar frequencies over time.

The two mechanisms of recombination and natural selection may also alter HCMV intrahost diversity. For other organisms, a positive correlation has been shown between recombination rate and diversity [25]. Both positive selection of beneficial alleles and negative selection against deleterious alleles will reduce diversity at a locus due to the fixation or clearance of mutations, respectively [26,27]. By uncoupling neutral variation from the beneficial or deleterious alleles, recombination can allow for higher levels of neutral variation to remain in the population. There is ample evidence of recombination in the HCMV genome [28-31], so this mechanism may play a role in shaping HCMV diversity. However, to date the recombination rate has not been quantified across the HCMV genome, while only a limited number of studies have analyzed the strength or frequency of selection targeting HCMV intrahost populations.

## Change of Populations in Human Hosts

With such high levels of intrahost diversity, researchers began to ask whether this leads to highly dynamic viral populations. For example, do the frequencies of specific mutations or genotypes change over time, and if so, how rapidly? Much research suggests that HCMV populations are generally stable over time. Longitudinal evolution studies of HCMV have predominantly focused on solid organ transplant recipients. In this patient population, genotyping via qPCR and sequencing of HCMV loci have shown mixed viral populations, but the composition of the mixed populations remains nearly constant in most patients over time [32-35]. A longitudinal study of congenitally infected infants showed similarly stable populations [22]. The predominant pattern from this work shows that HCMV populations contain a dominant sequence (which may represent > 90% of the total viral population) while the remaining population is occupied by a diverse array of largely stable sequences (i.e. minor variants) [15,16].

However, some caveats should be considered with this statement of longitudinal genetic stability. First, there may exist many thousands of stable mutations in an HCMV population [15,16,22,36] that do not rise in frequency over time, but importantly, are also not cleared from the population. These stable mutations (i.e., standing variation) provide a mechanism by which the virus could rapidly adapt to changing environments, such as new host compartments, immune pressure or antiviral drugs. Second, even if most mutations are stable, the change of only a few can dramatically alter viral fitness or pathogenesis. The increase in frequency of antiviral drug resistance mutations [37-39] clearly illustrates this point. Interestingly, antiviral treatment has also been shown to cause changes in non-drug resistance associated loci, such as gB, presumably due to linkage with a drug resistance mutation [33]. Lastly, HCMV populations that undergo spatial compartmentalization, as discussed in the next section, appear to undergo rapid evolution.

## Tissue compartmentalization of HCMV in Human Hosts

A hallmark of HCMV infections is dissemination to a wide range of host tissue compartments and cell types [40]. Due to the high levels of diversity associated with HCMV infections, researchers have begun to study whether viral diversity sampled from compartments differ, both in total level of variation and in specific genetic or genomic sequences observed. Indeed, early reports of diverse HCMV infections also demonstrated that HCMV sampled from different organs of healthy individuals could differ in gB genotype composition [14,41]. This finding has been supported in subsequent work studying other genomic loci [10,33,42]. Quantitative descriptions of the diversity have demonstrated significant differences in the level of diversity between compartments, with blood populations appearing more diverse than urine or intraocular populations [7,22]. However, the most striking example of compartmentalization may be differences in drug resistance genotypes observed in some patients. Several studies have shown that ganciclovir (GCV) resistance genotypes are not evenly distributed in patient compartments during treatment [43-46], with, for example, drug sensitive strains present in the cerebrospinal fluid and drug resistant strains in the blood [46].

Although the mechanism(s) explaining compartmentalization are not clear, there are at least two hypotheses that can be proposed. First, compartmentalization could result from the stochasticity of dissemination. Bottlenecking of the viral populations during dissemination will increase the proportion of mutations in the population that are governed by stochastic fluctuations in frequency (i.e., genetic drift) [47]. These stochastic changes can drastically skew the composition of populations in the distal relative to the original compartment [48]. Second, compartmentalization could result from natural selection during dissemination. Selection for mutants more fit in a compartment, for example due to cellular tropism, could lead to distinct populations in that tissue. Few studies have tested between these two models. There is evidence that selection [49] or both bottlenecks and selection can explain the observed compartmentalization [22] but the phenomenon needs to be studied in a wider range of patient populations to distinguish between these possibilities.

## HCMV Diversity, Evolution and Clinical Disease: Is there a connection?

We are now developing a clearer understanding of HCMV intrahost genetic diversity and evolution, but there is still the looming question of how these observations inform our understanding of HCMV disease. While this question is clinically relevant, work connecting diversity and pathogenesis is not fully developed, so discussion of its implications remains more speculative than others.

Many conflicting reports have examined the correlation between variation of HCMV genotype sequences and risk of transmission or disease progression and severity [41,50-54]. From this work, no clear conclusion can be reached about a possible effect of any single genotype on disease. However, subsequent research has shown that mixed genotype infections do correlate with increased viral load, HCMV disease severity, and even progression to AIDS in people with HIV infection [7,8,55,56]. Further, studies of intrauterine transmission of mothers with preconceptional immunity showed a higher rate of transmission and symptomatic congenital infections in mothers who were reinfected during pregnancy as compared to those who were not reinfected [18,19]. From these data, it is not clear if reinfection *per se* or merely the presence of multiple strains lead to the clinical outcomes. It has been demonstrated in a mouse model of cytomegalovirus infection that in vivo complementation occurs due to co-infection, allowing for the replication and dissemination of an attenuated strain due to transcomplementation from a wild type strain [57]. Thus, it is possible that multiple strain infections alter HCMV fitness by allowing a largely haploid virus to act as a polyploid.

Two specific areas where HCMV intrahost diversity may alter pathogenesis are dissemination and immune evasion. The possible role in dissemination is based on studies of the viral glycoproteins, proteins important in cell-to-cell spread and tissue tropism. In one report, many gB sequences were isolated from urine samples, but only a subtype of sequences were associated with plasma, suggesting that genetic diversity influences tissue tropism of the virus [41]. Similarly, variation in the gO protein alters the ratio of the endothelial-tropic gH/gL/UL128-131 complex to the fibroblast-tropic gH/gL/gO complex and thus was proposed to alter tropism and viral dissemination. From these data, it is possible to speculate that a diverse population of HCMV may harbor a subset of mutants that better replicate in distinct host compartments, and that viral diversity therefore enhances viral dissemination. In addition to dissemination, the possible effect of immune evasion on the relationship between viral intrahost diversity and pathogenesis has been suggested but not clearly proven. HCMV infections are regulated by both innate and adaptive immune responses. Variation in the *UL40* gene product modulates NK-cell mediated immunosurveillance by altering interaction of HLA-E with the CD94-NKG2A inhibitory receptor [58]. In studies of congenital infections, the intrahost diversity of *UL40,* particularly in the HLA-E binding domain, is significantly higher than the genome wide average [16]. Thus, mutants in the population likely display varied phenotypes in terms of NK-cell evasion, which may have profound effects on viral pathogenesis and the balance between productive and latent infections. Similar to NK cell evasion, variation in CD8^+^ T cell epitopes can alter the host adaptive immune response. Primary infection with mixed strains leads to the production of cross-reactive T cells, though exposure to some viral variants may limit the range of epitopes and viruses recognized by the T cell population [59]. Further, epitope recognition of the host T cell response has been shown to broaden over time in congenital infections [60], and thus may select against a wide range of viral mutations. Whether viral diversity is altered by and/or drives the adaptive immune response is unclear. Future work is needed to elucidate the directionality of this dynamic relationship between HCMV and its human host and the contribution of pre-existing and *de novo* viral mutants in the process.

## Conclusions and Future Directions

Here we highlight a few areas in which additional study could provide valuable information for ongoing and future efforts to lessen the burden of HCMV disease. An increasing body of evidence has demonstrated that HCMV is genetically diverse in human hosts, and there is tantalizing evidence that this diversity may contribute to pathogenesis. As such, HCMV intrahost diversity will remain as an important avenue to explore when developing novel antivirals and effective vaccines against this virus, as well as in designing treatment regimens - with previous results in other viruses suggesting that reduced diversity may indeed be associated with decreased pathogenesis [61,62]. Yet, in order for HCMV diversity to become a therapeutic target, it is necessary to better characterize the evolutionary processes responsible for this diversity. Relatedly, the need remains to better explore the observed correlative relationship between HCMV diversity and disease in order to disentangle cause from effect, and in so doing characterize the number of mutations important for the ultimate disease phenotype. Finally, relatively little attention has been given thus far to virus-host interactions, and further illuminating the role of variable host genetics as well as the co-evolutionary relationship between these organisms will likely become fruitful as sequencing costs continue to fall, thus enabling large scale paired human-HCMV sequencing duos. Encouraging, with these continued advances on the intersection of virology and population genetics, there is every reason to be optimistic that these aims will result in new tools to aid in our ongoing struggle with HCMV.

## Acknowledgments

We wish to thank our many colleagues who provided specimens from HCMV-infected patients that contributed to our understanding of intrahost evolution. This publication was supported by a grant from the National Institutes of Health (R01HD061959). The contents of this publication are solely the responsibility of the authors and do not necessarily represent the official views of the NIH.

